# DNA Methylation-Based Deconvolution Sheds Light on Glioblastoma Heterogeneity and Cell-Type Composition Associated with Patient Survival

**DOI:** 10.1101/2025.02.27.640603

**Authors:** Aviel Iluz, Nir Lavi, Hanna Charbit, Mijal Gutreiman, Masha Idelson, Debora Steiner, Etti Ben-Shushan, Aviad Zick, Amir Eden, Anat Mordechai, Moscovici Samuel, Yakov Fellig, Alexander Lossos, Joshua Moss, Benjamin E. Reubinoff, Iris Lavon

## Abstract

**Background:** IDH-wildtype glioblastoma (GBM) is an aggressive, heterogeneous brain tumor with limited treatment options. DNA methylation profiling allows detailed tumor characterization. This study applies methylation-based deconvolution to define GBM’s cellular composition and its association with patient outcomes.

**Methods:** We generated oligodendroglial precursor cells at various developmental stages from enriched human neural progenitor cultures and used their DNA methylation signatures, along with published signatures of brain tumor and tumor microenvironment–relevant cell types, to deconvolve 263 adult GBMs (Heidelberg-cohort). Tumor purity was estimated using RF_Purify. An independent cohort of 199 GBMs from TCGA and GEO, all treated with standard-of-care therapy, was similarly deconvolved, followed by Kaplan–Meier survival analysis to assess the prognostic value of neoplastic component proportions.

**Results:** Deconvolution uncovered distinct cellular compositions, consistent with single-cell RNA sequencing findings. Tumor purity analysis showed neoplastic fractions averaged 70% of tumor bulk, predominantly oligodendrocyte-like (43%), oligodendrocyte precursor-like (27%), astrocyte-like (19%), and mesenchymal stem cell-like (11%) populations. Non-neoplastic fractions were enriched for macrophages, vascular cells, and immune populations. A higher oligodendrocyte-like signature was linked to poorer survival (median survival 14.3 vs. 15.3 months; p = 0.017), while a higher astrocyte-like signature correlated with improved outcomes (15.3 vs. 13.4 months; p = 0.044). The astrocyte-to-oligodendrocyte ratio emerged as a strong prognostic marker, with a higher ratio predicting significantly longer survival (15.8 vs. 11.9 months; p < 0.00011).

**Conclusions:** Methylation-based deconvolution provides insight into GBM heterogeneity, highlighting the prognostic relevance of the astrocyte-to-oligodendrocyte ratio, which may guide personalized treatment strategies.

**Graphical Abstract:** A. We generated oligodendroglial precursor cells (OPs) at various developmental stages from enriched human neural progenitor cultures. A reference atlas of methylation signatures from 14 normal cell types, including the in vitro–generated cells, combined with published profiles of brain tumor and tumor microenvironment–relevant cell types, was applied to deconvolve 263 GBM samples based on their methylation profiles (Heidelberg cohort). Cell type proportions were analyzed with RF_Purify to estimate tumor purity and distinguish neoplastic from non-neoplastic components.
B. An independent cohort of 199 GBMs from TCGA and GEO, with available clinical data and all treated with standard-of-care therapy, was similarly deconvolved based on methylation profiles. Kaplan–Meier survival analysis assessed the prognostic impact of cell type proportions, identifying the astrocyte-to-oligodendrocyte ratio as the most significant marker. (Created with BioRender.com)

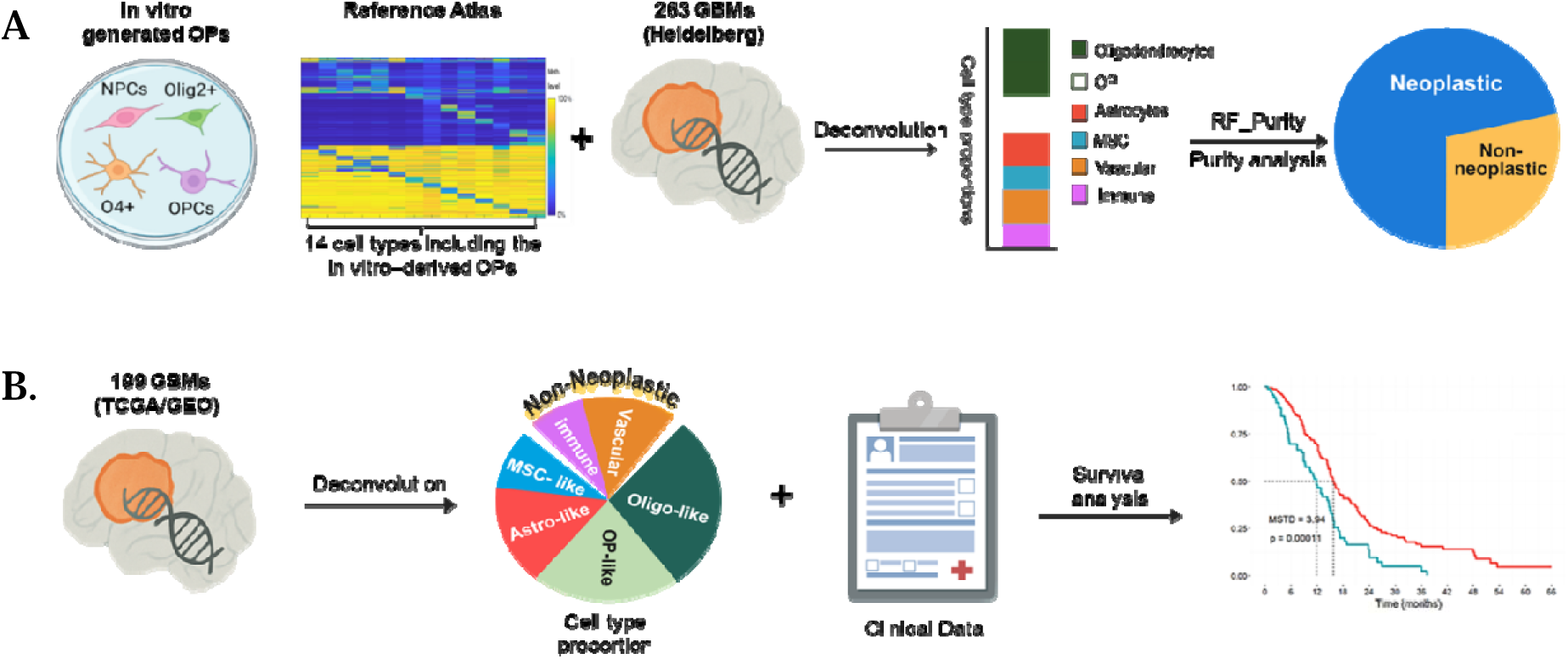

**Key points:**

- Methylation-based deconvolution incorporating in-vitro-derived oligodendrocyte precursor differentiation data reveals GBM heterogeneity
- GBM’s neoplastic fraction is dominated by oligodendrocyte-lineage cells
- Astrocyte/oligodendrocyte ratio is a strong prognostic marker for patient survival

**Importance of the study:** This study advances neuro-oncology research by using methylation-based deconvolution to uncover the cellular heterogeneity of glioblastoma. By constructing a comprehensive reference atlas and identifying cell-type-specific methylation signatures, it provides prognostic insights into the distinct cellular compositions of glioblastoma. Combining deconvolution results with purity analysis enables the differentiation between neoplastic and non-neoplastic tumor components. This approach complements single-cell RNA sequencing while offering greater clinical applicability, as DNA methylation profiling can be performed on FFPE or fresh frozen samples, unlike the more costly and tissue-sensitive single-cell methods. Importantly, the identification of the astrocyte-to-oligodendrocyte ratio as a strong prognostic marker highlights key cellular determinants of glioblastoma survival. These findings deepen our understanding of glioblastoma biology and offer a practical tool for patient stratification.

## Introduction

IDH-wildtype glioblastoma (GBM), is an aggressive and lethal brain cancer, known for its rapid growth and resistance to standard treatments.^1,2^ A key feature of GBM is its extensive cellular heterogeneity, which presents challenges for treatment. The tumor contains a complex mix of neoplastic and non-neoplastic cells, including glioma-initiating cells, immune cells, endothelial cells, and glial cells.^2-5^

Genome-wide DNA methylation profiling has become a transformative tool for classifying Central Nervous System (CNS) tumors, enabling patient stratification beyond traditional diagnostics.^6,7^ In GBM, beyond confirming the diagnosis, DNA methylation profiling helps identify distinct subclasses, including receptor tyrosine kinase I (RTK1), receptor tyrosine kinase II (RTK2), and mesenchymal (MES).^6-8^ Studies of newly diagnosed GBMs reveal significant spatial diversity within methylation subclasses, underscoring both the complexity of tumor classification and the critical role of intratumoral heterogeneity in driving treatment resistance and disease recurrence.^9^

Understanding the cellular drivers of GBM heterogeneity is essential for advancing research and developing effective therapies. Various methods, including proteomic and transcriptomic analyses, have been used to explore this complexity.^2-4,10^ Recent breakthroughs in single-cell RNA sequencing (scRNA-seq) have provided new insights into the cellular diversity of GBM, revealing distinct subpopulations of neoplastic cells and their interactions with the tumor microenvironment (TME).^2-5^ A landmark study^3^ identified four distinct cellular states within the neoplastic fraction of GBM: astrocyte-like (AC-like), oligodendrocyte progenitor cell-like (OPC-like), mesenchymal-like (MES-like), and neural progenitor cell-like (NPC-like). These findings underscore the tumor’s complex differentiation and plasticity.

Analyzing DNA methylation signatures offers an alternative approach to investigating GBM cellular composition, other than scRNA-seq.^11^ Each cell type has a unique methylation signature that reflects its tissue identity.^12,13^ It is well-established that tumors exhibit stable methylation signatures, even after oncogenic events, enabling precise sub-classification of brain tumors.^14,15^ These signatures have been validated in thousands of CNS tumors, facilitating accurate diagnoses and better treatment guidance.^6,16-18^ This method was incorporated into the 2021 WHO Classification of CNS tumors.^18^

The DNA methylation signature of a tumor reflects the cumulative methylation patterns of all its constituent cell types, including both normal and neoplastic cells. This composite profile captures the cellular admixture within the tumor, providing insights into its complex cellular makeup.^11^ Methylation-based deconvolution using a reference atlas of normal cell-type methylation signatures estimates tissue composition and has been applied to analyze tissue contributions in circulating cell-free DNA (cfDNA).^19,20^ A similar deconvolution methodology has been applied to investigate the composition of non-neoplastic immune cells within the GBM TME.^11^

To the best of our knowledge, methylation signatures have not been specifically used to investigate the neoplastic components of GBM. In our study we take a comprehensive approach to analyze both neoplastic and non-neoplastic components of GBM tumors using methylation-based deconvolution and methylation-based purity analysis. By analyzing methylation signatures from 462 GBM samples across three datasets, we aim to elucidate GBM heterogeneity encompassing all relevant cell types within the tumor microenvironment, with particular focus on the neoplastic fraction, and to investigate associations between the abundance of specific cell types and patient survival.

This analysis provides valuable insights into the cell of origin of GBM, a critical factor in understanding early tumorigenic processes and identifying potential targets for early intervention. Since DNA methylation profiles preserve signatures of the originating cell type, they offer a reliable method for tracing cellular origins. When combined with our uniquely developed in vitro-derived human neural progenitors and oligodendrocyte-lineage precursors methylation dataset, this approach allows for a comprehensive exploration of the cell of origin in GBM.^21^

To validate and contextualize our findings, we compared our results with data obtained through complementary methodologies, including scRNA-seq and the deconvolution of immune cell profiles from bulk glioma DNA methylation data.^3,11,22-24^ This integrative approach is expected to enhance our understanding of GBM’s cellular complexity and inform future prognostic and therapeutic strategies.

## Materials and Methods

### Generation of human NP and Oligodendrocyte Lineage Precursor Cells

Highly enriched cultures of neural progenitors were derived from human embryonic stem cells (hESCs) using an established protocol (designated in this study as "NP").^25^ Briefly, hESCs (HAD-C100^26^) at passages 22 to 28 were cultured on human recombinant Laminin 521 (Biolamina, Sundbyberg, Sweden) in NutriStem medium supplemented with growth factors (Biological Industries, Beit Haemek, Israel) and 50 units/mL penicillin and 50 μg/mL streptomycin (Pen-Strep, Thermo Fisher Scientific, Waltham, MA, USA). Oligodendroglial precursor cells at various developmental stages were derived from these enriched NP cultures as previously described.^25^ Briefly, NP were first guided to differentiate into OLIG2-positive progenitor cells (designated as "OLIG2") and subsequently directed toward the oligodendroglial lineage. This process resulted in an enriched population of Olig2+/Nkx2.2+ oligodendrocyte progenitor cells (designated as "OPC"), which were further differentiated into O4+ pre-oligodendrocytes (designated as "pre-oligodendrocytes").

### Immunofluorescent Staining of hESC-Derived Oligodendroglial Lineage Cells

To characterize the developmental stages of oligodendroglial lineage cells, markers representing different maturation steps were detected by immunofluorescence. Briefly, cells were plated on glass coverslips pretreated with poly-D-lysine (30–70 kDa, 10 mg/ml) and laminin (4 mg/ml; both from Sigma, St. Louis, MO) and cultured for 3–5 days. The cells were then fixed with 4% paraformaldehyde and incubated with mouse monoclonal anti-Olig2 (1:150; EMD Millipore, Burlington, MA) and rabbit anti-Nkx2.2 (1:50; Novus Biologicals, Centennial, CO) antibodies. Alexa Fluor 555-conjugated donkey anti-mouse antibody (1:150; Invitrogen/Thermo Fisher Scientific, Waltham, MA) and Alexa Fluor 488-conjugated donkey anti-rabbit antibody (1:150; Jackson ImmunoResearch Laboratories, West Grove, PA) were used for detection.

For the pre-oligodendrocyte stage, cells were incubated with mouse anti-O4 antibody (1:150; R&D Systems, Minneapolis, MN), followed by fixation with 4% paraformaldehyde and incubation with Alexa Fluor 488-conjugated donkey anti-mouse antibody (1:150; Jackson ImmunoResearch). Nuclei were counterstained with 4′,6-diamidino-2-phenylindole (DAPI; Vector Laboratories, Burlingame, CA). Specimens were visualized using an Olympus BX61 fluorescence microscope (Olympus, Hamburg, Germany).

### Flow Cytometry

Human ESC-derived neural progenitor cells (NPCs) were immunostained with mouse anti-PSA-NCAM-PE (1:50) and mouse anti-A2B5-APC (1:50) antibodies, or the corresponding isotype controls (all from Miltenyi Biotec, Waltham, MA). For each sample, at least 10^4 cells were analyzed on a CytoFLEX flow cytometer (Beckman Coulter, Indianapolis, IN).

### DNA methylation profiling

Genome-wide DNA methylation profiling of NP, OLIG2, OPC, and pre-oligodendrocytes was performed using the Illumina EPIC array, with data generated at the Genomics and Proteomics Core Facility at the German Cancer Research Center (DKFZ) in Heidelberg, Germany. Data processing, filtering, and normalization were carried out using the *minfi* R package (v1.34.0).

### Data collection

We collected 450K or EPIC Illumina cytosine-guanine dinucleotides (CpG) methylation signature data to assemble the various sample sets analyzed in this study, as detailed below. All data were processed as beta-value CSV files. To ensure consistency, EPIC beta-value probe set were intersected with the 450K beta-value probe set, retaining only common probes for downstream analysis.

### Tumor sample sets for deconvolution and survival analysis

For the deconvolution analysis, we selected DNA methylation profiles of GBM samples classified as "Glioblastoma, IDH-wildtype" according to the WHO 2016 classification of CNS tumors,^17^ obtained from the Heidelberg reference cohort (GSE90496). To focus on adult GBM, we included 263 samples further sub-classified into receptor tyrosine kinase I (RTK1; n=64), receptor tyrosine kinase II (RTK2; n=143), or mesenchymal (MES; n=56) subtypes, as defined by Capper et al.^6^ This classification was originally derived using the DNA methylation-based brain tumor classifier (version 11.4), which improves diagnostic accuracy and standardization.^6^

Methylation data of glioma-initiating cell (GIC) sample set (n=20) was obtained from a publicly available dataset referenced in Vinel et al.^27^ (GSE155985). In this study, the authors isolated GICs from GBM samples, which were classified by methylation profiling into RTK1, RTK2, and MES.

For survival analysis, and further validation of our results we obtained methylation data of an independent cohort of 199 GBM cases with available clinical data from multiple sources: (1) a cohort of 84 TCGA samples diagnosed as primary GBM,^8^ and (2) a cohort of 115 GBM samples from two GEO datasets (53 from GSE60274,^28^ 62 from GSE195640^29^). We selected only patients that were treated according to the Stupp protocol, which consists of radiation therapy with concurrent and adjuvant temozolomide. All the 199 GBM patients were successfully subclassified into the RTK1 (n=39), RTK2 (n=97), and MES (n=63) subclasses using the DKFZ brain tumor classifier (v12.8) on the Heidelberg Epignostix Classifier platform (https://app.epignostix.com).

Methylation profiles of non-glial tumor sets were obtained from the TCGA database and represent the average methylation profiles of samples from four tumor types: bladder urothelial carcinoma (BLCA) (n=292), breast carcinoma (n=721), kidney renal papillary cell carcinoma (KIRP) (n=210), and prostate adenocarcinoma (PRAD) (n=127).

Methylation data (CpG beta-value CSV files) for various normal cell types, including 25 signatures from Moss et al.,^20^ microglia,^11^ MSCs (GSM4077441–GSM4077443, GSM4078810–GSM4078818), astrocytes (GSM3938231), oligodendrocytes,^19,20^ and cortical neurons (GSE98203) were sourced from multiple databases and studies.

### Bioinformatics and statistical analysis

#### Constructing reference atlases

We constructed reference atlases by assembling methylation profiles of the relevant cell types for each atlas. To identify tissue-specific CpG sites within each atlas, we applied the feature selection method described by Moss et al.^20^ Briefly, CpGs (based on hg19 coordinates) with low variance (<0.1%) across the methylation atlas or missing values were excluded. Methylation values for each CpG across cell types were normalized by their sum, and the top 100 hypermethylated CpGs per cell type were selected based on specificity. A similar procedure was applied to the reversed methylation matrix to identify hypomethylated CpGs. For each cell type, both the top 100 hypermethylated and hypomethylated CpGs, along with neighboring CpGs within 50 bp, were included in the reference matrix. To further refine the feature set, pairwise-specific CpGs were iteratively selected by projecting the atlas onto the current CpG set, calculating Euclidean distances between cell types, and adding CpGs that best distinguished the most similar pairs at each iteration.

#### GBM tumors reference atlases

We used two reference atlases for GBM deconvolution analysis. The initial reference atlas included 16 cell type signatures: B-cells, CD4 T-cells, CD8 T-cells, NK cells, neutrophils, vascular endothelial cells, monocytes, microglia, MSC, cortical neurons, astrocytes, oligodendrocytes, NP, OLIG2, OPC, and pre-oligodendrocytes, covering a total of 4,712 CpG sites. Based on the results from deconvolving Heidelberg set of 263 GBM samples, we refined this atlas to include 14 cell type signatures by merging OLIG2, OPC, and pre-oligodendrocytes into a single oligodendrocyte-lineage precursor component (designated as "OP"), reducing the total to 4,111 CpG sites. This refined atlas was subsequently used to deconvolve Heidelberg set GBM, TCGA and GEO GBM, and GIC samples.

### Deconvolution of Methylation Signatures Using NNLS Linear Regression

Non-negative least squares (NNLS) linear regression was employed, as described by Moss et al.,^20^ to deconvolve the GBM methylation signature beta-value matrices into cell-type components, using the relevant reference atlas.

### Validation of the Deconvolution method

To validate the ability of the methylation-based deconvolution method to correctly identify expected cell type signatures, we applied it to methylation data from non-GBM sources using a custom reference atlas we constructed, comprising cell type signatures relevant to the tested samples, as follows:

#### Deconvolution of non-glial tumors

We applied methylation-based deconvolution to non-glial tumors, including bladder urothelial carcinoma (BLCA), breast carcinoma, kidney renal papillary cell carcinoma (KIRP), and prostate adenocarcinoma (PRAD). The reference atlas used for this analysis comprised 25 cell type signatures, sourced from publicly available data by Moss et al.^20^ and spanning 7,390 CpG sites. The results showed that the predominant cell type in each tumor corresponded to the expected tissue of origin: bladder cells (74%) in BLCA, breast cells (62%) in breast cancer, kidney cells (79%) in KIRP, and prostate cells (90%) in PRAD (Figure S1).

#### Deconvolution of normal cell types

We applied methylation-based deconvolution to 11 samples representing various normal cell types. The reference atlas used in this analysis included 11 cell type signatures derived from publicly available data by Moss et al.^20^, using an N-1 approach: for each cell type, one corresponding sample was excluded from the atlas for validation and subsequently deconvolved using the remaining data. The atlas comprised a total of 3,274 CpG sites. The analysis showed that over 90% of the estimated cell type proportions in each sample matched the expected normal cell type. For example, the cortical neuron sample was estimated to contain 100% cortical neurons (Figure S2).

### Methylation-based tumor purity

Tumor purity was calculated using the *RF_purify* R package (v0.1.2) with the ABSOLUTE method. This method analyzes Illumina methylation data to estimate purity by applying a Random Forest machine learning algorithm, trained on copy number variations (CNVs) inferred from methylation patterns.^30^

### Survival analysis

Kaplan–Meier analysis and log-rank tests were conducted on 199 GBM samples using the *survminer* (v0.5.0) and *survival* (v3.7-0) R packages.^31^ We assessed overall survival (OS) differences based on the median proportion of each cell type across all samples. Patients with proportions greater than the median were classified as "high", while those with proportions less than the median were classified as "low", and their survival outcomes were compared. Astrocytes and Oligodendrocytes showed significant overall survival differences with opposite survival trends (higher overall survival for higher proportions versus lower overall survival for higher proportions). To assess the prognostic significance of the ratio between astrocyte-like and oligodendrocyte-like signatures within the neoplastic fraction, we determined the optimal cutoff for the astrocyte-to-oligodendrocyte ratio. The optimal cutoff was established by evaluating the ratio values across the entire range, with the threshold identified as the point yielding the most significant log-rank test result. This cutoff dichotomized the ratios into ‘high’ and ‘low’ astrocyte-to-oligodendrocyte ratio groups, which exhibited distinct survival probabilities, consistent with the previous methodology.^32^

To validate this cutoff and address potential overfitting, we applied a permutation-based approach. The Astrocyte-to-Oligodendrocyte ratio was randomly shuffled across patients (100K iterations) while maintaining survival times and censoring status, and log-rank p-values were recomputed for each permutation. The permutation p-value was calculated as the proportion of permuted p-values less than the observed p-value, with statistical significance defined at p<0.05. A low permutation p-value indicates that the observed p-value is unlikely under randomized conditions, reflecting the robustness of the selected cutoff. This approach confirms the cutoff’s reliability and significance, distinguishing it from effects attributable to chance or dataset-specific overfitting.

### Statistical Analysis

To assess significant differences in cell type proportions across GBM subclasses, pairwise Student’s two-tailed t-tests were conducted for each subclass pair. Two-tailed t-tests were also used to evaluate the statistical significance of Pearson correlation coefficients between cell type proportions and purity scores. T-test were performed using R, Python or Microsoft Excel provided built-in t-test functions. For survival analysis, the log-rank test was applied as previously described.

To compare the expected cell-type proportion fractions between the survival analysis GBM set (n=199) and the observed results from the initial GBM set (n=263), we performed a chi-square goodness-of-fit test using the *chisq.test* function in R (with parameters *simulate.p.value = TRUE, B = 2000)*.

For comparisons of subclass distributions between patient survival probability groups, we used the chi-square test for independence with the *chisq.test* function in R.

A significance threshold of 0.05 was used for all statistical tests.

### Graph and Plots

All graphs and plots were generated using R (v4.0.3+), Python 3.11, Microsoft Excel, or BioeRender.com.

## Results

### Generation of Neural Progenitors and Human Oligodendrocyte Lineage Precursor Cells

Neural progenitors (NP) were successfully derived from human embryonic stem cells and subsequently differentiated into oligodendroglial precursor cells through a staged process (Figure 1). These NP cultures (Figure 1A) were guided to generate OLIG2-positive progenitor cells (OLIG2) (Figure 1B), which were further directed toward the oligodendroglial lineage. This resulted in enriched populations of Olig2+/Nkx2.2+ oligodendrocyte progenitor cells (OPCs) (Figure 1C) and their subsequent differentiation into O4+ pre-oligodendrocytes (pre-oligodendrocytes) (Figure 1D), confirming progression along the expected developmental trajectory.

**Figure 1:**
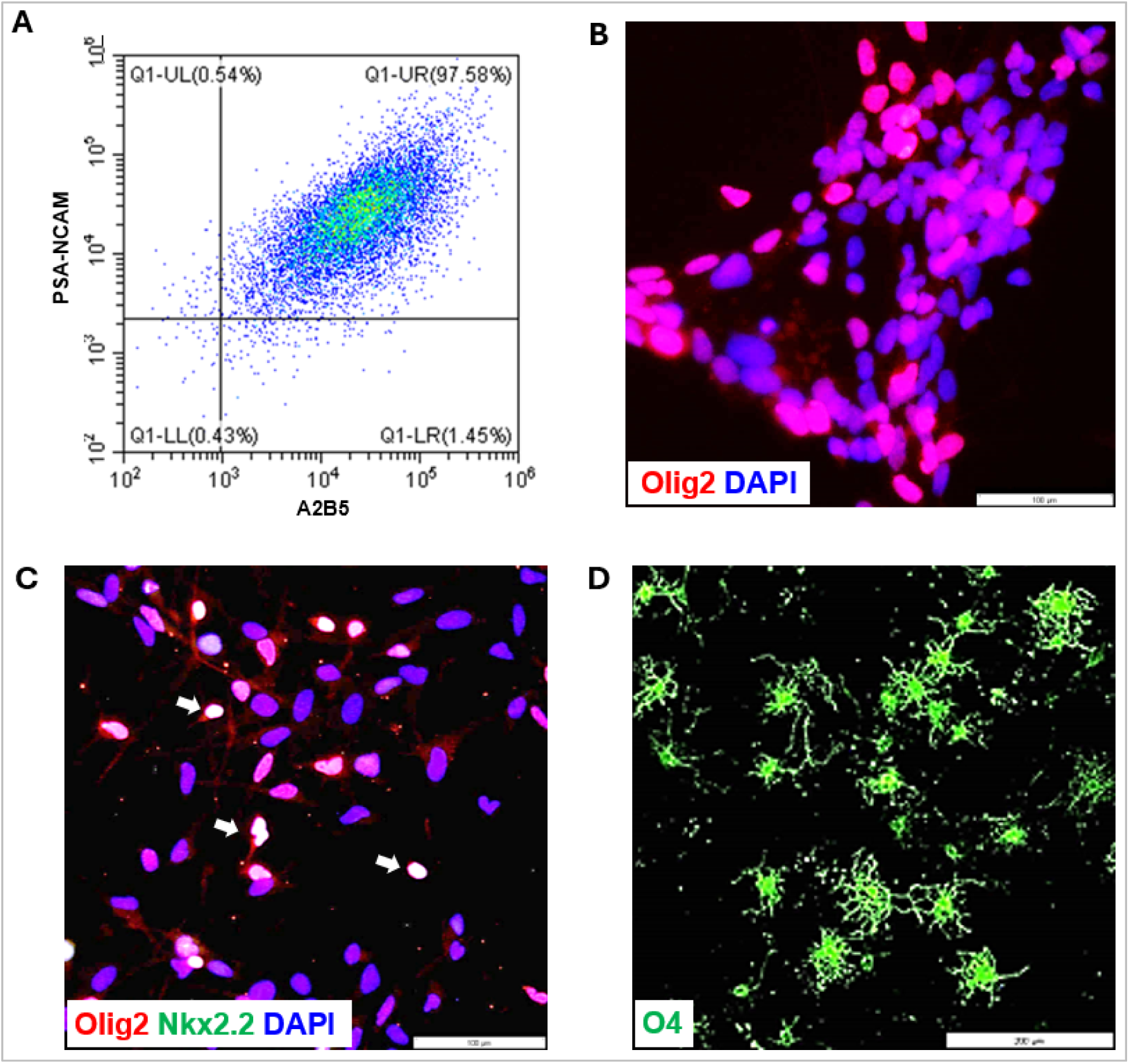
Differentiation of hESCs towards oligodendroglial lineage cells. (A) Dot plot presentation of NPs co-expressing PSA-NCAM and A2B5; (B-D) Immunostaining showing the expression of specific markers, characterizing different stages of the development of oligodendroglial lineage cells. (B) OLIG2-positive progenitors, (C) OPCs co-expressing Olig2 and Nkx2.2 (marked by white arrows); (D) pre-oligodendrocytes expressing O4. Scale bars: (B-C) 100mm, (D) 200mm.

### Identification of Cell Type Composition and Proportions in GBM and Its Subtypes: RTK1, RTK2, and MES

To investigate cell type composition in GBM, we constructed a reference atlas of DNA methylation signatures critical for the deconvolution of GBM samples and quantification of cell type proportions. This atlas encompassed methylation data from 16 distinct cell types, derived from our previously published data^19,20^ and several publicly available datasets, as detailed below. It included cortical neurons, astrocytes, oligodendrocytes, and various immune cells (B cells, CD4L T cells, NK cells, CD8L T cells, and neutrophils), as well as tumor-associated macrophages (TAM; monocytes and microglia^11^), and mesenchymal stem cells (MSCs). To account for the potential differentiation states of cancer cells, we further extended the atlas by incorporating methylation profiles from our in vitro-generated neural progenitors and three defined stages within the oligodendrocyte lineage. These stages represent a continuum of differentiation and include OLIG2-positive progenitors (characterized by expression of the OLIG2 transcription factor essential for oligodendrocyte development), oligodendrocyte progenitor cells (OPCs), and pre-oligodendrocytes identified by O4 expression. This atlas was generated by selecting distinct differentially methylated cytosine-guanine dinucleotides (CpGs) for each cell type, following previously described and validated methodologies.^20^ In total, it comprises 4,712 CpG sites across these cell types (**Figure S3**).

To estimate the cellular composition of bulk GBM tumor samples, we applied a deconvolution method adapted from cfDNA tissue-origin studies.^20^ Using 450K methylation data we analyzed 263 adult GBM samples, classified according to their methylation profiles as previously described.^6^

In GBM samples glial cells constituted the majority (62%) of the cellular composition, including contributions from oligodendrocytes-like signature (26.3%), astrocytes-like (13.4%), and pre-oligodendrocytes-like (O4+, 22%). Cortical neurons signature was minimally represented (2.6%). Additionally, GBM displayed 7.3% mesenchymal stem cells-like (MSC), 7.8% TAM comprising microglia (3.6%) and monocytes (4.2%), 13% vascular endothelial cells and 7.2% immune cells (B-cells, CD4 T-cells, NK cells, CD8 T-cells, and neutrophils). Neural progenitors showed effectively no contribution (**Figure S4**).

To further refine the analysis of GBM cell type composition, OLIG2, OPC, and pre-oligodendrocytes were merged into a single component termed oligodendrocyte-lineage precursor (OP). This adjustment was necessary because OLIG2 and OPC contributed minimally to GBM proportions (average 0.3 ± 1.2%). However, their integration ensured representation while preserving the accuracy of cell type contribution. The final reference atlas included 14 cell-types (**Figure 2**).

**Figure 2.**
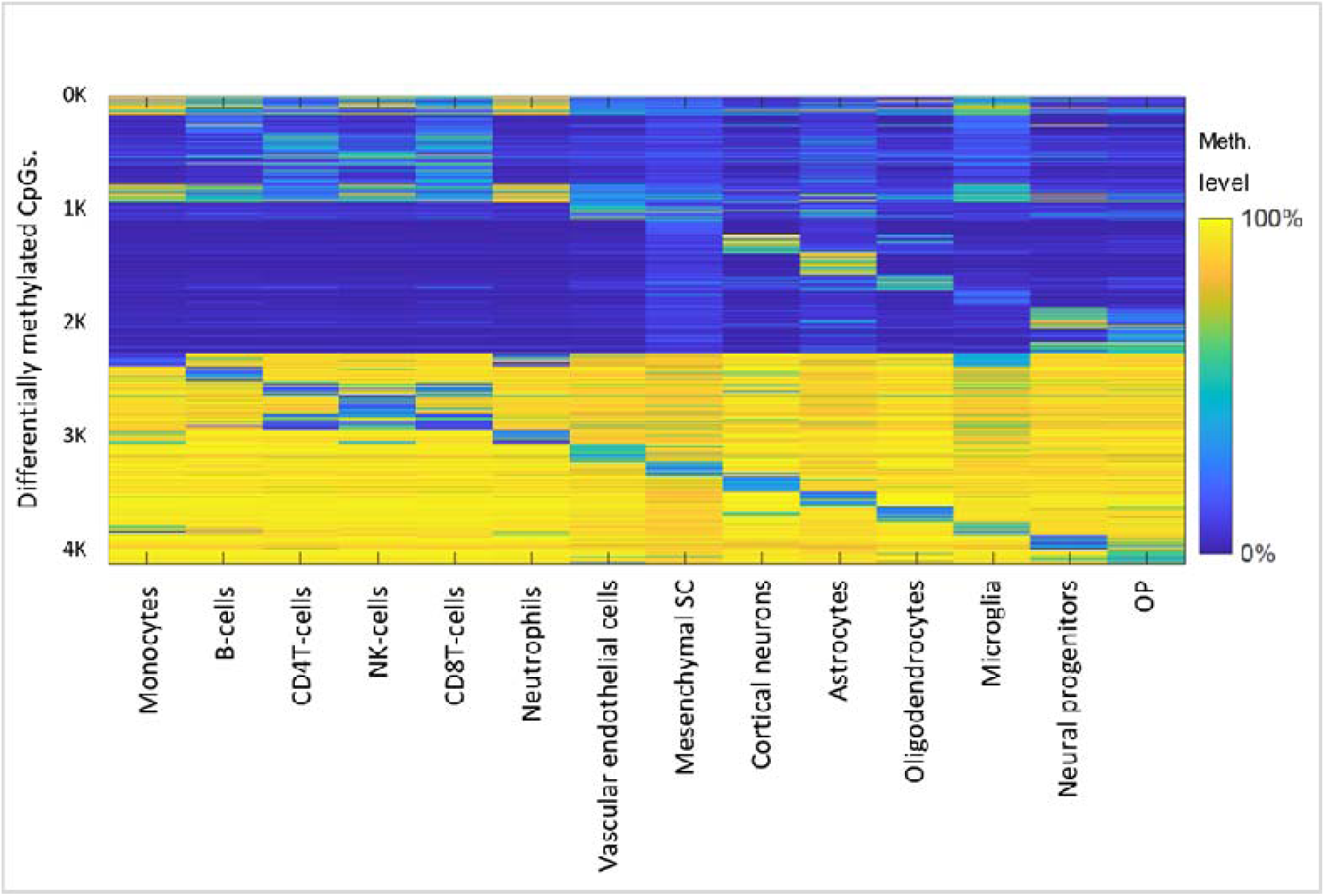
Reference atlas for GBM deconvolution and cell-type identification using methylation signatures. The methylation atlas comprises 14 tissues/cell types (columns) across 4111 CpGs sites (rows) that are located in 2610 genomic blocks of 500 bp each. Feature selection was carried out as described in a previous study.^20^ Specifically, the top 100 uniquely hypermethylated CpGs and hypomethylated CpGs for each cell type, yielding a total of 2800 tissue-specific CpGs, with neighboring CpGs within 50 bp were also included.

This adjustment had minimal impact on the estimated proportions of astrocytes, cortical neurons, MSCs, TAMs, vascular endothelial cells, and immune cells. However, a slight difference emerged in the oligodendrocyte lineage, with the proportion of oligodendrocytes increasing from 26.3% to 29.5%, while pre-oligodendrocytes, which accounted for 22% in the separated analysis, were reduced to 18.9% in the combined OP component. (**Figures 3A, S4**).

**Figure 3.**
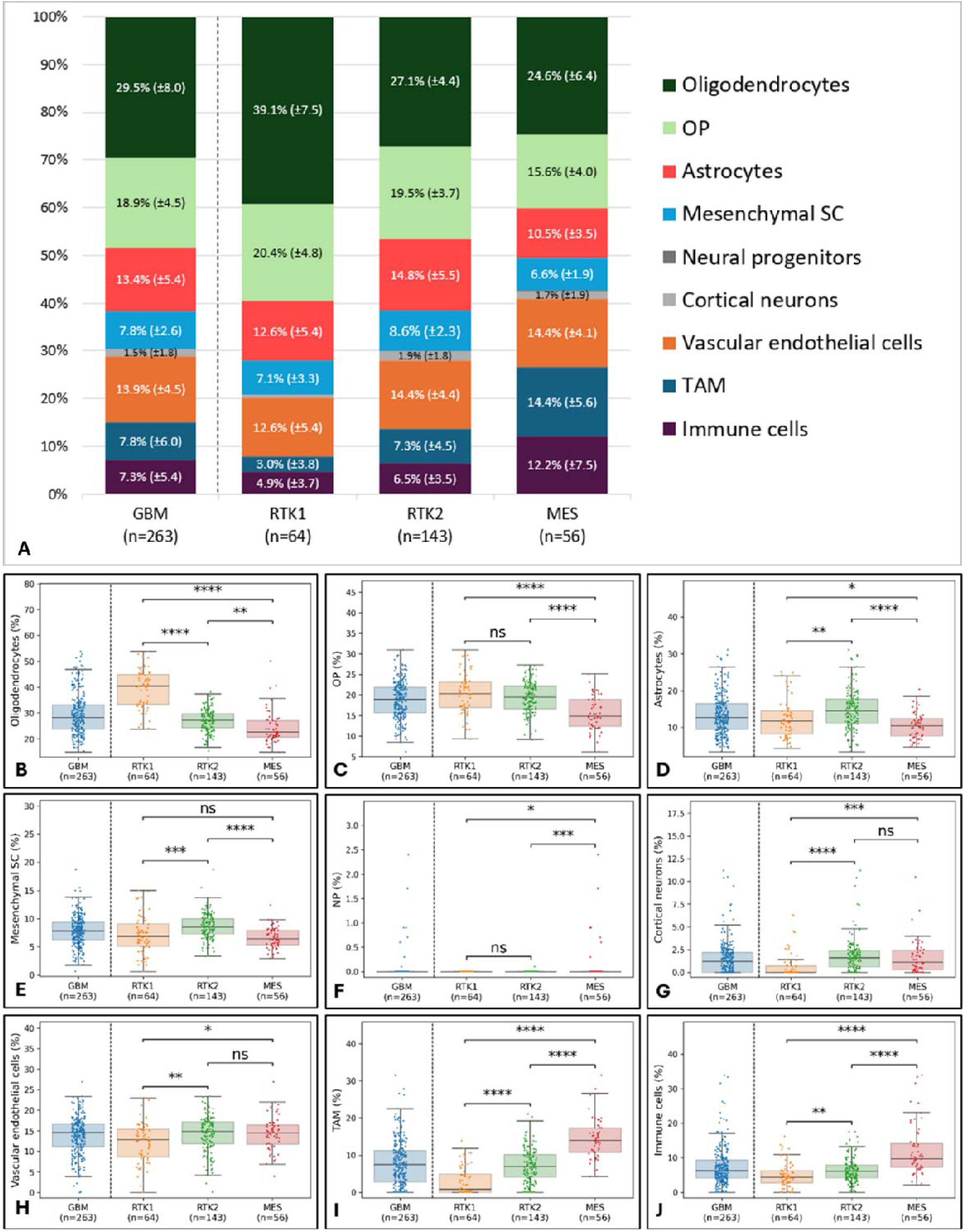
Cell-type proportions derived from deconvolution analysis in GBM and their differences across GBM subtypes, based on atlas of 14 cell types. (A) Bar plot summarizing the average proportions of 14 cell types (combined into 9 groups) from the deconvolution of methylation profiles of 263 samples shown in A-I (leftmost bar), subdivided into RTK1 (n=64), RTK2 (n=143), and MES (n=56). The average percentage and standard deviation for each cell type are displayed within its respective bar. (B-J) Boxplots depicting the proportions of 9 cell-types or cell type groups derived from the deconvolution of methylation data across GBM samples (n=263, blue, leftmost), subdivided into RTK1 (n=64, orange), RTK2 (n=143, green), and MES (n=56, pink). The 9 panels correspond to (B) oligodendrocytes, (c) oligodendrocyte-lineage precursors (OP), (D) astrocytes, (E) mesenchymal stem cells (MSC), (F) neural progenitors, (G) cortical neurons, (H) vascular endothelial cells, (I) tumor-associated macrophages (TAM; combined microglia and monocytes), and (J) immune cells (combined B cells, CD4 T cells, NK cells, CD8 T cells, and neutrophils). The central line represents the median, box edges indicate the interquartile range (IQR), and whiskers extend to 1.5 times the IQR. Black lines denote significant differences between groups, determined by pairwise two-tailed Student’s t-test. Exact p-values are reported in the main text. (*p<0.05; **p<0.01; ****p<0.0001; ns: not significant).

Furthermore, our findings revealed significant variation in cell type proportions among GBM subclasses. Specifically, astrocyte proportions differed significantly between RTK1 and RTK2 (12.6% vs. 14.8%, p<0.01) and between RTK2 and MES (14.8% vs. 10.5%, p<1.60E-07), but not between RTK1 and MES (**Figure 3D**). Oligodendrocyte proportions also showed significant pairwise differences in all comparisons: RTK1 vs. RTK2 (39% vs. 27%, p<1.012E-32), RTK1 vs. MES (39% vs. 24.6%, p<2.39E-20), and RTK2 vs. MES (27% vs. 24.6%, p<1.85E-03) (**Figure 3B)**. Additionally, significant differences were observed in the proportions of other cell types, including OP, MSC, immune, and TAM cell populations. All p-values were calculated using a two-tailed t-test **(Figures 3B-J)**.

### Assessing the Neoplastic and Non-Neoplastic Components of GBM Using RF_Purify Method

Using cell type proportions from the deconvolution analysis, we aimed to distinguish between neoplastic and non-neoplastic components in GBM tumors. This was done using the RF_Purify method, which applies the ABSOLUTE score to infer tumor purity.^33^ We correlated purity scores with the proportions of seven cell types identified in the deconvolution analysis, excluding cortical neurons (average 2%) and NP (average 0%) due to their minimal contribution. The analyzed components included immune cells, TAM, vascular endothelial cells, MSC, astrocytes, OP, and oligodendrocytes (**Figure 4**).

**Figure 4.**
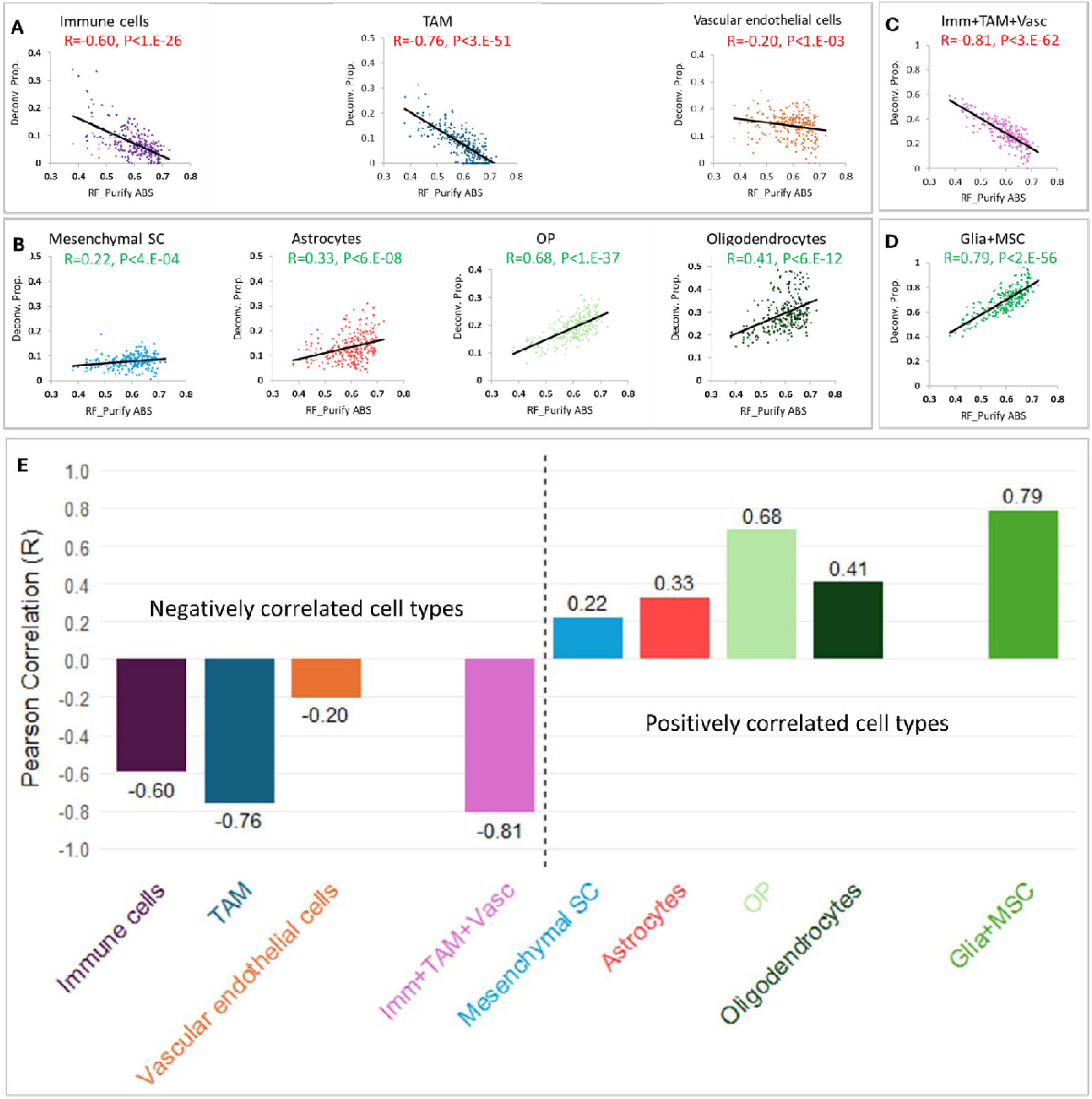
Correlation between cell-type proportions and purity estimates calculated by RF_Purify_ABSOLUTE. (A-D) Correlation between cell type proportions (Y axis) derived from the deconvolution of all 263 GBM samples and sample purity (X axis) estimated by the RF_Purify algorithm using the ABSOLUTE method. The analyzed cell types include (A) total immune cells (purple), tumor-associated macrophages (TAM; dark blue), and vascular endothelial cells (dark orange); (B) mesenchymal stem cells (MSC; blue), astrocytes (red), oligodendrocyte-lineage precursors (OP; light green), and oligodendrocytes (dark green); (C) the combined total of the three negatively correlated cell types (immune cells + TAM + vascular endothelial cells; light purple); (D) the combined total of the four positively correlated cell types (three glial cell types + MSC; green). (E) An overview histogram illustrates the Pearson correlation (R) values (Y axis) between proportions for nine cellular components showed in A-D (X-axis) and sample purity. The dashed line divides the graph into two sections, with the far-right bar in each section representing the correlation for the combined positively (glia + MSC, green) or negatively (immune cells + TAM + vascular endothelial cells, pink) correlated cell types.

Significant negative correlations were observed between tumor purity and the proportions of immune cells (R=-0.60, p<1.E-26), TAM (R=-0.76, p<3.E-51), and vascular endothelial cells (R=-0.20, p<1.E-03) (**Figures 4A, 4E**). In contrast, tumor purity correlated positively with glial lineage cells, including astrocytes (R=0.33, p<6.E-08), OP (R=0.68, p<1.E-37), oligodendrocytes (R=0.41, p<6.E-12), and with MSC (R=0.22, p<4.E-04) (**Figures 4B, 4E**). When the proportions of all four positively correlated components (three glial lineage cell types and MSC) were combined additively as a single component, the positive correlation with tumor purity was stronger (R=0.79, p<2.E-56), suggesting these components collectively contribute substantially to the neoplastic portion of the tumor (**Figures 4D, 4E**). On the other hand, when the proportions of all negative correlated components (immune cells, TAM, and vascular endothelial cells) were combined additively as a single component, the negative correlation with tumor purity was stronger (R=-0.81, p<3.E-62), suggesting these components collectively contribute substantially to the non-neoplastic normal portion of the tumor (**Figures 4C, 4E**). All p-values were calculated using a two-tailed correlation t-test.

### Comparison of Methylation-Based Deconvolution Results with scRNA-Seq Studies

Our study found that the neoplastic fraction, estimated from 263 GBM samples using methylation-based deconvolution, averaged 70% (±11%), demonstrating strong concordance with prior estimates derived primarily from scRNA-seq^3,10,22-24^, which reported an average of 72% (range 52%–87%) (Table 1, Figure 5). Overall, the tumor purity levels are highly consistent across methods. While most cell type proportions were broadly comparable to earlier findings, a few notable differences emerged. For instance, vascular endothelial cells were markedly higher in our analysis, averaging 13.9% (±4.5%) compared to 1.7% (range 0.3%–2.8%) in prior studies. We also observed a slightly higher average of immune cells (7.3% ±5.4%) relative to the 3.3% reported elsewhere. These discrepancies may reflect differences in sample collection, tissue processing, or methodological biases, particularly the tendency of certain cell types to be underrepresented in single-cell RNA-seq due to fragility or dissociation inefficiency. Despite these variations, our findings reinforce the robustness of methylation-based deconvolution in capturing the major cellular components of GBM.^10,11,22-24^

**Figure 5:**
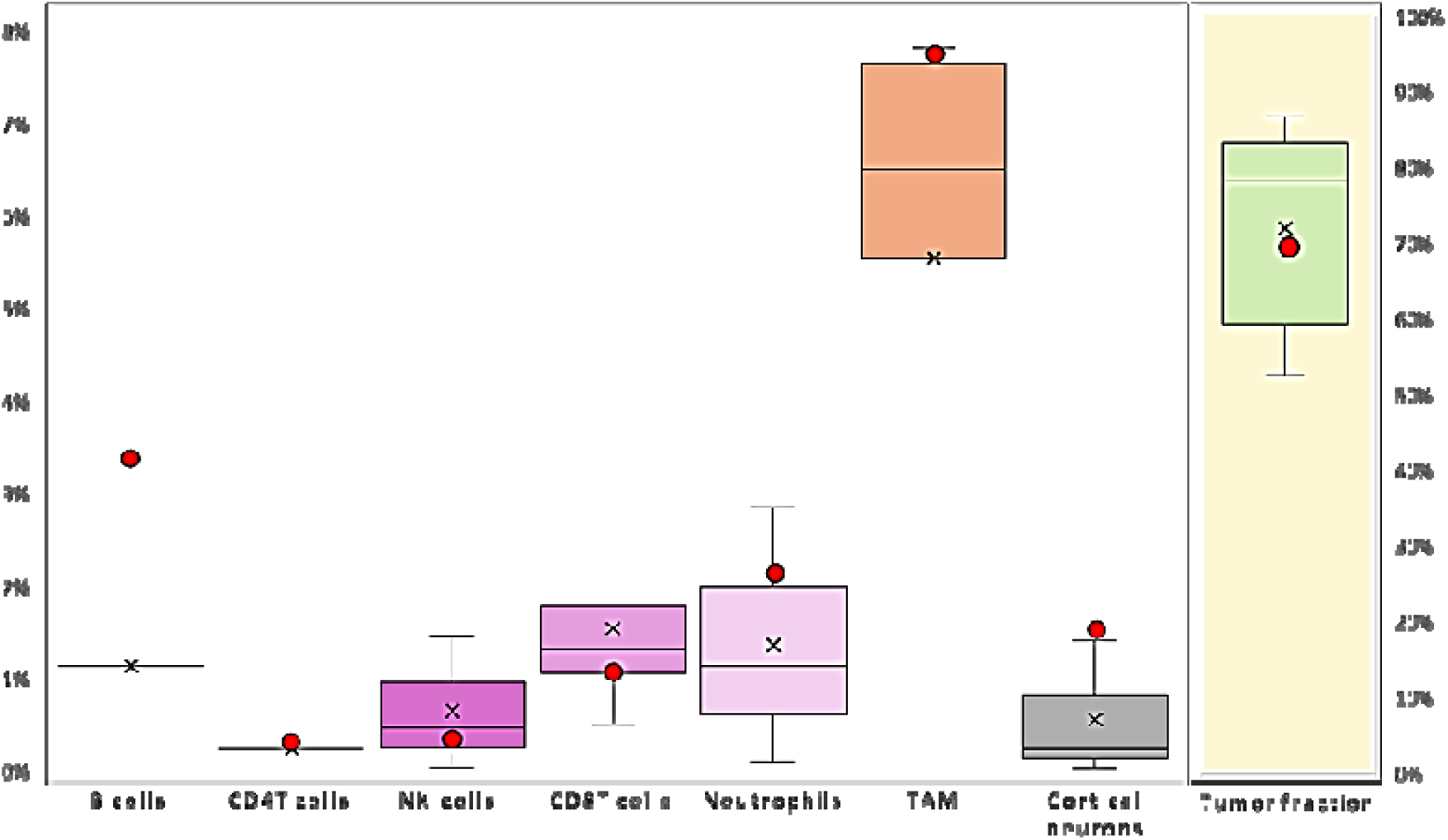
Comparison of proportions of different cell types between our study and other studies. Boxplots illustrating the distribution of cell type proportions and the neoplastic fraction in GBM samples from other studies (as detailed in Table 1), with each boxplot representing a specific cell type. The boxplot on the far right depicts the neoplastic fraction on a 100% scale. Our findings are indicated by a red circle for comparison. The central line in each boxplot shows the median, the X marks the mean, the box edges indicate the interquartile range (IQR), and the whiskers extend to 1.5 times the IQR.

**Table 1:** Comparison of Methylation-Based Deconvolution Results with other studies.

| TME fractions | cell types | Neftel | Yu | Darmanis | Yuan | Singh | other studies (avg.) | our study (avg.) |
| --- | --- | --- | --- | --- | --- | --- | --- | --- |
| **Non-Neoplastic fraction** | B cells | - | - | - | - | 1.1% | 1.1% | 3.4% |
|  | CD4T cells | - | - | - | - | 0.2% | 0.2% | 0.3% |
|  | NK-cells | 0.0% | 1.4% | - | - | 0.4% | 0.6% | 0.4% |
|  | CD8T cells | 1.2% | 3.0% | - | 0.5% | 1.3% | 1.5% | 1.1% |
|  | Neutrophils | 0.1% | 2.8% | - | - | 1.1% | 1.3% | 2.2% |
|  | TAM | 5.4% | 6.4% | 7.7% | 0.2% | 7.5% | 5.4% | 7.8% |
|  | Cortical neurons | - | - | 1.4% | 0.0% | 0.2% | 0.5% | 1.5% |
|  | Normal Oligodendrocytes | 2.8% | 9.1% | 6.9% | 1.6% | 6.2% | 5.3% | - |
|  | Normal Astrocytes | - | 0.6% | 1.4% | - | - | 1.0% | - |
| **Neoplastic fraction** |  | 87% | 52% | 59% | 83% | 78% | 72% | 70% |

Summary of the proportions of various cell types and neoplastic fraction identified in our study (the right most column) compared to those reported in other studies and their average. The left four data columns represent the scRNA-Seq-based studies: Neftel et al.<sup>3</sup>, Yu et al.,<sup>24</sup> Darmanis et al.<sup>22</sup> and Yuan et al.<sup>23</sup> The fifth column on the right displays findings from methylation-based analysis by Singh et al.<sup>11</sup> The two rightmost columns display the averages from other studies and our study.

### Validation of Purity Analysis: Deconvolution Results in Bulk GBM Tumors Versus Glioma-Initiating Cells

Our purity analysis identified three glial cell types and mesenchymal stem cells as contributing to the neoplastic fraction of GBM, in agreement with previous single-cell RNA-seq studies. To further validate this finding, we applied our deconvolution approach to DNA methylation profiles of CD133 (Prominin-1)-enriched glioma-initiating cells (GICs), which represent the neoplastic compartment of the tumor. These profiles were obtained from a previously published study^27^ and derived from GBM tumors classified as RTK1, RTK2, and MES (see Methods). This comparison allowed us to evaluate whether the cell-type signals associated with the neoplastic fraction in bulk tumors are retained in a highly purified population of tumor cells.

As expected, deconvolution of GICs revealed a significantly higher tumor fraction composed of three glial lineage cell types and MSCs compared to GBM bulk tumors, with 88% in GICs versus 70% in GBM (p<2.2E-20, two-tailed t-test, **Figure S5**). Specifically, GICs exhibited significantly increased proportions of astrocytes (19% vs. 13%, p<8.7E-04), OP (26% vs. 19%, p<3.1E-05), and MSCs (16% vs. 8%, p<5.6E-11), while oligodendrocyte proportions remained similar (28% vs. 30%, p<0.21). In contrast, vascular endothelial cells, immune cells, and TAMs were significantly reduced in GICs compared to GBM bulk tumors (vascular endothelial cells: 7% vs. 14%, p<2.4E-09; immune cells: 3% vs. 7%, p<1.1E-07; TAMs: 0% vs. 8%, p<6.5E-54) (**Figure S5**). All -values were calculated using a two-tailed t-test. These findings demonstrate that cell types positively correlated with tumor purity in GBM (**Figure 4B**) are more abundant in GICs, whereas those negatively correlated with tumor purity (**Figure 4A**) are present at lower proportions.

### Correlation Between Cell Type Proportions and Patient Survival: Prognostic Significance of the Astrocyte-to-Oligodendrocyte Ratio

To evaluate whether the abundance of specific cell types is associated with patient survival, we analyzed 199 GBM samples with available methylation clinical data from TCGA and GEO. Using our methylation-based deconvolution approach, we estimated cell type proportions in this cohort and confirmed consistency with the original set of 263 samples (chi-square goodness-of-fit test, *p* = 0.96; Figure S6). We then assessed the association between cell type proportions and overall survival using Kaplan–Meier analysis (Figure 6).

**Figure 6:**
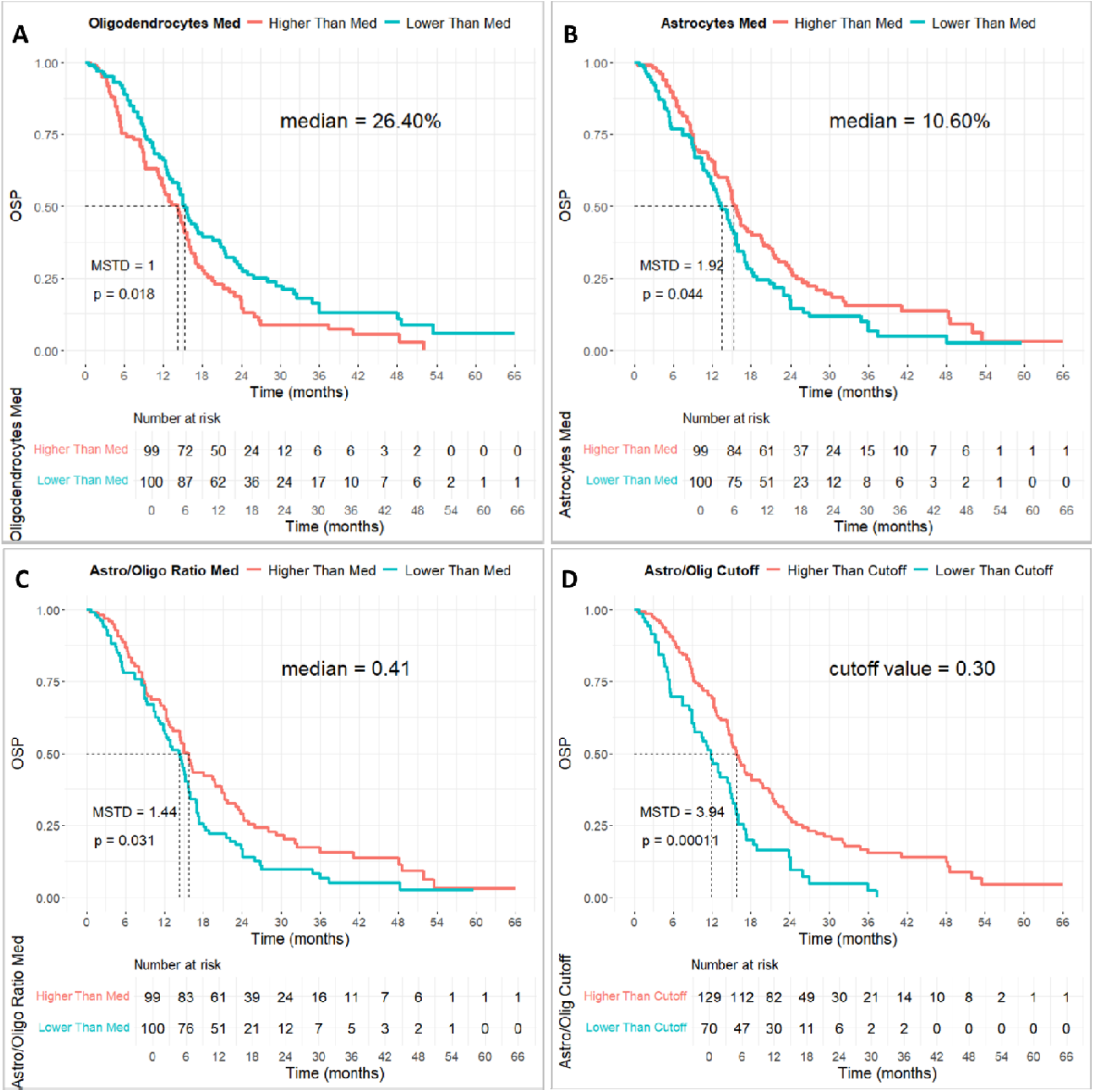
Kaplan-Meier plots based on cell type proportions. Kaplan-Meier survival analysis of 199 cases showing significant differences in overall survival probability (OSP) based on cell type proportions. The median survival time difference (MSTD) between the groups is indicated in months. (A-C) Groups are stratified into high and low groups (High > median, Low < median) according the cell type proportion. (A) Oligodendrocyte, (B) Astrocyte, (C) Astrocyte-to-oligodendrocyte proportion ratio. (D) Astrocyte-to-oligodendrocyte proportion ratio, divided into high and low groups using a cutoff value (0.3) determined by the threshold yielding the most significant log-rank test result, validated by a permutation-based approach. (MSTD, median survival time difference; p-values based on log-rank test).

We found that oligodendrocyte proportions lower than the median (26.4%) was significantly associated with higher overall survival probability (OSP), whereas higher proportions were linked to lower OSP (median survival time [MST]: 15.3 vs. 14.3 months; median survival time difference [MSTD]: 1 month; p < 0.018, log-rank test; Figure 6A).

In contrast, higher astrocyte proportions (above the median, 10.6%) were associated with improved OS, while lower proportions correlated with shorter OSP (MST: 15.3 vs. 13.4 months; MSTD: 1.9 months; p<0.044, log-rank test; **Figure 6B).** A similar trend was observed for microglia, where proportions above the median (3.2%) correlated with increased OSP compared to lower proportions (MST: 15.7 vs. 14.3 months; MSTD: 1.4 months; p<0.007, log-rank test; **Figure S7B**). No significant correlations were found for other cell types or cell type groups (immune cells, TAM, vascular endothelial cells, OP, and MSC) (**Figure S7A-F**). Building on the contrasting effects of astrocyte and oligodendrocyte abundance on OS, and considering previous studies highlighting dynamic shifts in cellular states within the GBMs’neoplastic fraction,^2-4,34^ we investigated the impact of the astrocytes to oligodendrocytes (Astro/Olig) ratio on survival. The median Astro/Olig ratio effectively startified the petients into high- and low-survival groups. A ratio above the median (>0.41) was significantly associated with increased OSP (MST = 15.7 months), whereas a lower ratio corresponded to reduced OSP (MST = 14.3 months), with an MSTD of 1.44 months (p<0.03, log-rank test, **Figure 6C**). To evaluate the Astro/Olig ratio as a prognostic biomarker, we determined the optimal cutoff that best separated the survival groups (see Methods). A cutoff of 0.3, corresponding to an Astro/Oligo ratio of 1:3.3, was identified. Patients with a ratio above this threshold exhibited significantly improved OSP (MST = 15.8 months), those with a lower ratio had a reduced OSP (MST = 11.9 months), with an MSTD of 3.9 months (p<0.00011, log-rank test, **Figure 6D**).

Notably, the average overall survival differed by 6 months between the ‘high’ and ‘low’ Astro/Oligo ratio cutoff groups. The higher ratio group had an average overall survival of 18.1 months, while the lower ratio group had an average overall survival of 11.6 months.

To validate this cutoff and mitigate potential overfitting, we applied a permutation-based approach with 100K random iterations on the ratio data (see Methods). The permutation p-value was 0.0001, representing the proportion of permuted p-values ≤ the observed p-value (0.00011). This validation underscores the robustness of the cutoff, differentiating it from random effects or dataset-specific overfitting.

Given the impact of the Astro/Oligo ratio on overall survival and the differences in astrocyte and oligodendrocyte proportions observed across the three GBM subclasses, RTK1, RTK2, and MES (**Figure 3**), we examined their association with survival outcomes to determine whether the observed survival differences were driven by a specific subclass.

We found that patients with an Astro/Olig ratio above the defined cutoff, which was associated with long-term survival (LTS), were predominantly from the RTK2 subclass (64%), whereas only 7% belonged to the RTK1 subclass. In contrast, patients with an Astro/Olig ratio below the cutoff, linked to short-term survival (STS), had a higher proportion of the RTK1 subclass (43%) compared to the RTK2 subclass (20%). Unlike the RTK1 and RTK2 subclasses, which exhibited opposite distributions between LTS and STS groups, the MES subclass did not follow this pattern (**Figure S8**).

However, a survival analysis based solely on the GBM subclass did not reveal significant differences in overall prognosis (**Figure S7G**).

## Discussion

Our study establishes methylation-based deconvolution as a powerful tool for unraveling GBM’s cellular composition, shedding light on tumor heterogeneity and linking cell type proportions to patient survival.

By leveraging a reference atlas of methylation profiles from GBM microenvironment cell types, particularly through the inclusion of in vitro-differentiated human oligodendrocyte-lineage precursors, we successfully delineated the neoplastic cell composition of GBM using DNA methylation-based deconvolution. To our knowledge, this is the first demonstration of neoplastic cell type composition in GBM revealed through methylation-based deconvolution, a result previously demonstrated only with techniques such as RNA-seq. Additionally, our approach enabled the identification of key cellular components within the tumor microenvironment.

Notably, glial cells dominated the GBM composition (62%), primarily oligodendrocytes-like, oligodendrocyte precursor-like and astrocytes-like signatures, while cortical neurons contributed minimally. Additionally, we identified presence of mesenchymal stem cells-like (MSC), vascular endothelial cells, tumor-associated macrophages (TAM), and immune cells, further underscoring the complexity of the GBM microenvironment.

Our results were consistent with prior analyses,^3,10,11,22-24^ with the notable exception of a higher vascular endothelial cell proportion (13%), consistent with a recent report indicating 9.9% vascularization in GBM.^35^ This finding underscores the well-documented angiogenic nature of GBM^36^ and reinforces the reliability of our approach in capturing GBM’s vascular complexity.

To distinguish neoplastic from non-neoplastic components, we used the RF_Purify method, which leverages the ABSOLUTE score to infer tumor purity. This analysis confirmed that oligodendrocyte-lineage precursors, astrocytes, oligodendrocytes, and MSCs, constitute the neoplastic fraction, while immune cells, TAM, and vascular endothelial cells predominantly belong to the tumor microenvironment. The aggregated neoplastic fraction averaged 70%, consistent with prior estimates (∼72%).^3,10,11,22-24^

Aligned with prior research,^3,35^ our findings indicate that astrocytes-like and oligodendrocytes-like signatures constitute a significant part of the GBM neoplastic fraction. The consistently reported low abundance of normal astrocytes and oligodendrocytes in GBM samples^3,10,11,22-24^ supports the conclusion that elevated proportions we detected primarily reflect the neoplastic component.

We propose that our astrocyte and MSC methylation signatures correspond to the AC-like and MES-like states, respectively, which were identified by scRNA-seq^3-5^ as part of the neoplastic fraction of GBM. Our analysis indicates that early oligodendrocyte lineage stages (OLIG2L or OPC-like cells) and neural progenitors contribute minimally to GBM composition. Instead, most oligodendrocyte-lineage cells display advanced differentiation, as reflected by pre-oligodendrocyte (O4-positive) and mature oligodendrocyte methylation signatures. Assigning the methylation signature of oligodendrocyte-lineage precursors (OP) and mature oligodendrocytes to the less differentiated NPC-like and OPC-like transcriptional signatures reported in scRNA-seq^3-5^ as part of the neoplastic fraction remains challenging. The discrepancy between our findings and the previous studies^3-5^ likely stems from methodological differences between single-cell transcriptional profiling and bulk methylation analysis. Further investigations, particularly single-cell methylation studies, are needed to validate this interpretation. Despite these methodological differences, both approaches support the notion that these populations represent distinct stages within the oligodendrocyte differentiation lineages.

We demonstrate that the oligodendrocyte differentiation lineage accounts for approximately 50% of the bulk tumor and a striking 71% of the neoplastic fraction. These findings extend prior work by Liu et al. ^37^, Persson et al ^38^, and Ligon et al.^39^, which showed that oligodendrocyte precursor cells can give rise to GBM under defined oncogenic conditions. By revealing that oligodendrocyte-like and oligodendrocyte precursor-like cells comprise 71% of the neoplastic fraction, our results strongly reinforce the involvement of the oligodendrocyte lineage in GBM pathogenesis and suggest that pre-oligodendrocytes may serve as cells of origin for GBM.

To evaluate the accuracy of our neoplastic cell composition estimates, we applied DNA methylation-based deconvolution to CD133-enriched tumor dissociates from 20 GBM patients.^40^ As expected, the neoplastic fraction was higher than in bulk tumors (∼90% vs. ∼70%), consistent with enrichment for cancer cells. The remaining ∼10% consisted largely of endothelial cells (∼6.8%), likely reflecting GBM’s prominent vasculature and minor contamination from tumor-associated blood vessels during CD133+ cell enrichment. Significant differences in cell-type proportions were observed among GBM subclasses RTK1, RTK2, and MES, including variations in TAM and immune cell fractions (e.g., B cells, CD4+ T cells, NK cells, CD8+ T cells, and neutrophils). These findings are consistent with transcriptomic and immunohistochemistry analyses, which report increased tumor-infiltrating lymphocytes and TAM in the Mesenchymal GBM subtype compared to non-Mesenchymal subtypes.^11,41-43^ Our results extend these subclassifications by revealing distinct differences in cellular composition detected through methylation-based deconvolution.

A key finding of our study is the significant association between GBM cellular composition and patient survival. Higher proportions of astrocytes were linked to improved overall survival, while increased proportions of oligodendrocytes and OPs were associated with poorer outcomes. These results align with previous studies showing that proliferative potential is highest in OPC- and NPC-like tumor states, and lowest in astrocyte-like (AC-like) states.^3,4^

Furthermore, we identified the astrocyte-to-oligodendrocyte ratio as a key prognostic biomarker. This aligns with the findings that the AC-like state has reduced tumor-initiating potential compared to oligodendrocyte-lineage components, which exhibit greater tumorigenicity in mouse models.^4^

Although RTK1, RTK2, and MES tumors exhibited distinct cellular compositions, these differences did not translate into significant overall survival variations across subtypes, consistent with findings from other studies.^41,44^. Instead, as suggested above, survival differences were driven by the astrocyte-to-oligodendrocyte ratio, suggesting that intertumoral cellular heterogeneity may be a more powerful prognostic indicator than subclassification alone.

Nevertheless, patients with lower survival times, as determined by a higher astrocyte-to-oligodendrocyte ratio, had a greater proportion of GBM tumors classified as RTK2 relative to RTK1, and vice versa. Since RTK2 showed higher proportions of astrocytes and lower proportions of oligodendrocytes, compared to RTK1, this may explain the survival differences between the two subclasses.

In conclusion, DNA methylation-based deconvolution offers a robust framework for resolving the cellular composition of GBM. We identify the astrocyte-to-oligodendrocyte ratio as a prognostic biomarker linked to patient survival. This approach provides clinically meaningful insights from initial tumor resections and stands out as faster, more cost-effective, and broadly applicable than single-cell RNA sequencing, compatible even with FFPE and archived samples. By avoiding tissue dissociation, it supports routine clinical implementation, with the potential to improve patient stratification, guide personalized therapy, and ultimately impact survival outcomes.

## Limitations of the Study

This study has several limitations. First, the DNA methylation profiles of in vitro-differentiated neural progenitors and oligodendrocyte-lineage precursors may not fully capture the methylation landscapes present in GBM tumors, potentially affecting the accuracy of our findings. Second, the survival analysis was conducted on a cohort predominantly receiving standard-of-care therapy, which may limit its applicability to patients undergoing alternative treatments or to underrepresented GBM subtypes. Further research is required to evaluate the broader applicability of these results.

## Required statements

### Ethics

**All human subject data used in this study were obtained from online databases, including The Cancer Genome Atlas (TCGA) and GEO.**

### Author Contributions

Conceptualization, A.I., A.E., J.M, I.L.; methodology, A.I., N.L., J.M, BE. R, M.I, D.S, E. BS, I.L. MS; software, A.I., N.L, J.M.; validation, A.I., J.M. M.I, A.Z. I.L.; formal analysis, A.I., I.L.; investigation, A.I, I.L.; resources, A.I., H.C, M.G., M.I, D.S., EB.S, M.S, J.M, BE, R, I.L.; data curation, A.I., N.L., J.M. BE. R, M.I, D.S; writing—original draft preparation, A.I., I.L.; writing-review and editing, A.I., N.L, J.M., I.L., A.L, A.M; Y.F. visualization, A.I., N.L., J.M., A.L, A.M, I.L.; supervision, I.L.; project administration, I.L.; funding acquisition, I.L. All authors have read and agreed to the published version of the manuscript.

### Data Availability

**Reference atlases used for deconvolution analysis and clinical data for the survival analysis cohort are available in the supplementary data.**

## Acknowledgments

The results shown here are in whole or part based upon data generated by the TCGA Research Network: https://www.cancer.gov/tcga.

We sincerely thank Andreas von Deimling and his laboratory for conducting the EPIC methylation profiling of the in vitro differentiated neural progenitors^45^ and oligodendrocyte precursor (OP) cells, which greatly contributed to this research.

We also extend our heartfelt thanks to Leslie and Michael Gaffin for their generous support.

