## Supplementary figures for "DNA Methylation-Based Deconvolution Sheds Light on Glioblastoma Heterogeneity and Cell-Type Composition Associated with Patient Survival"

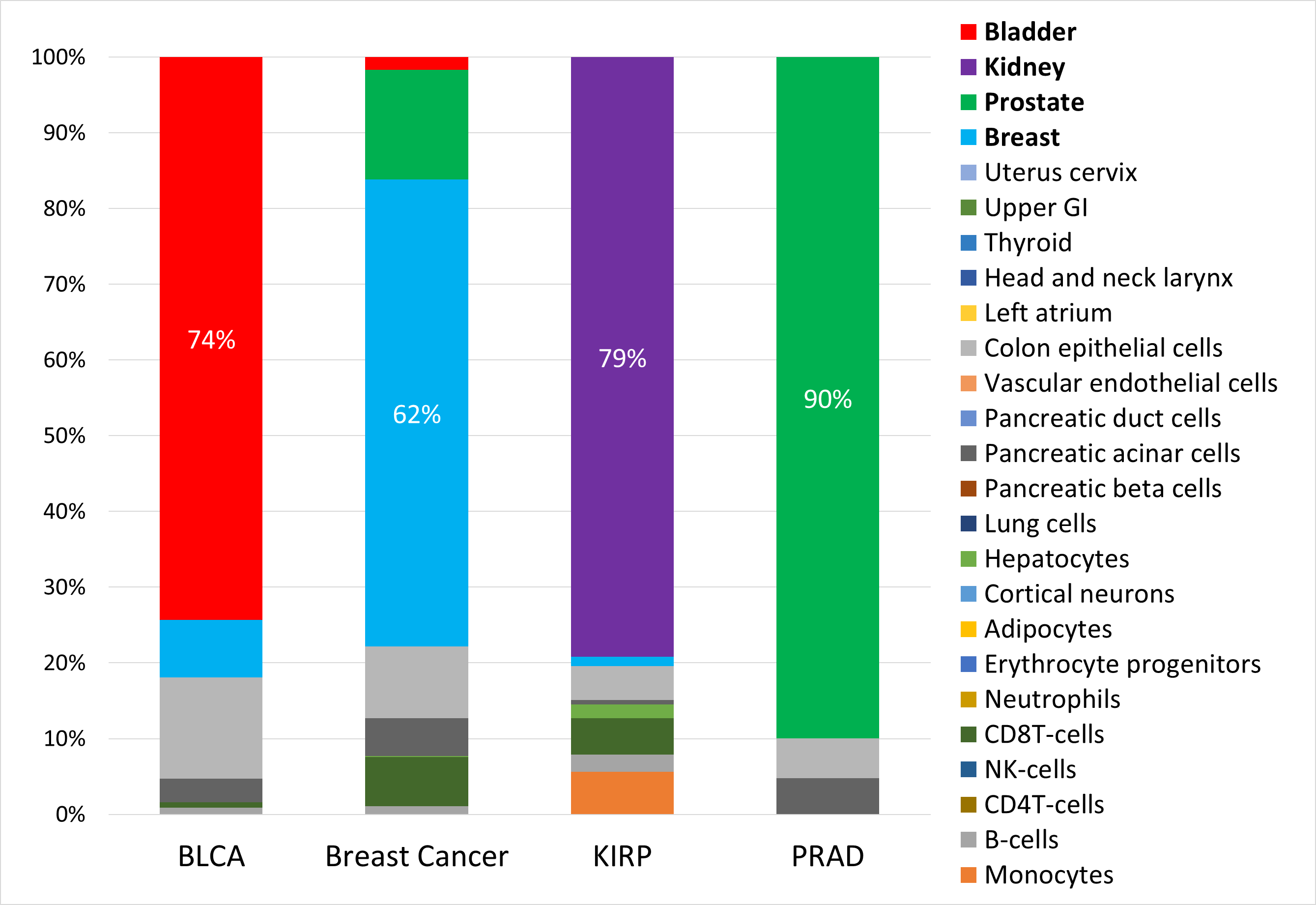

Figure S1. Methylation-based deconvolution of non-glial tumors.

Bar plots comparing the average proportions of normal cell types derived from the deconvolution of methylation profiles from four different non-glial tumors (bladder urothelial carcinoma (BLCA), breast cancer, kidney renal papillary cell carcinoma (KIRP) and prostate adenocarcinoma (PRAD). Deconvolution was performed using a 25-cell-type reference atlas from Moss et al.^20^ The specified percentage indicates only the highest cell type proportion in each tumor type.

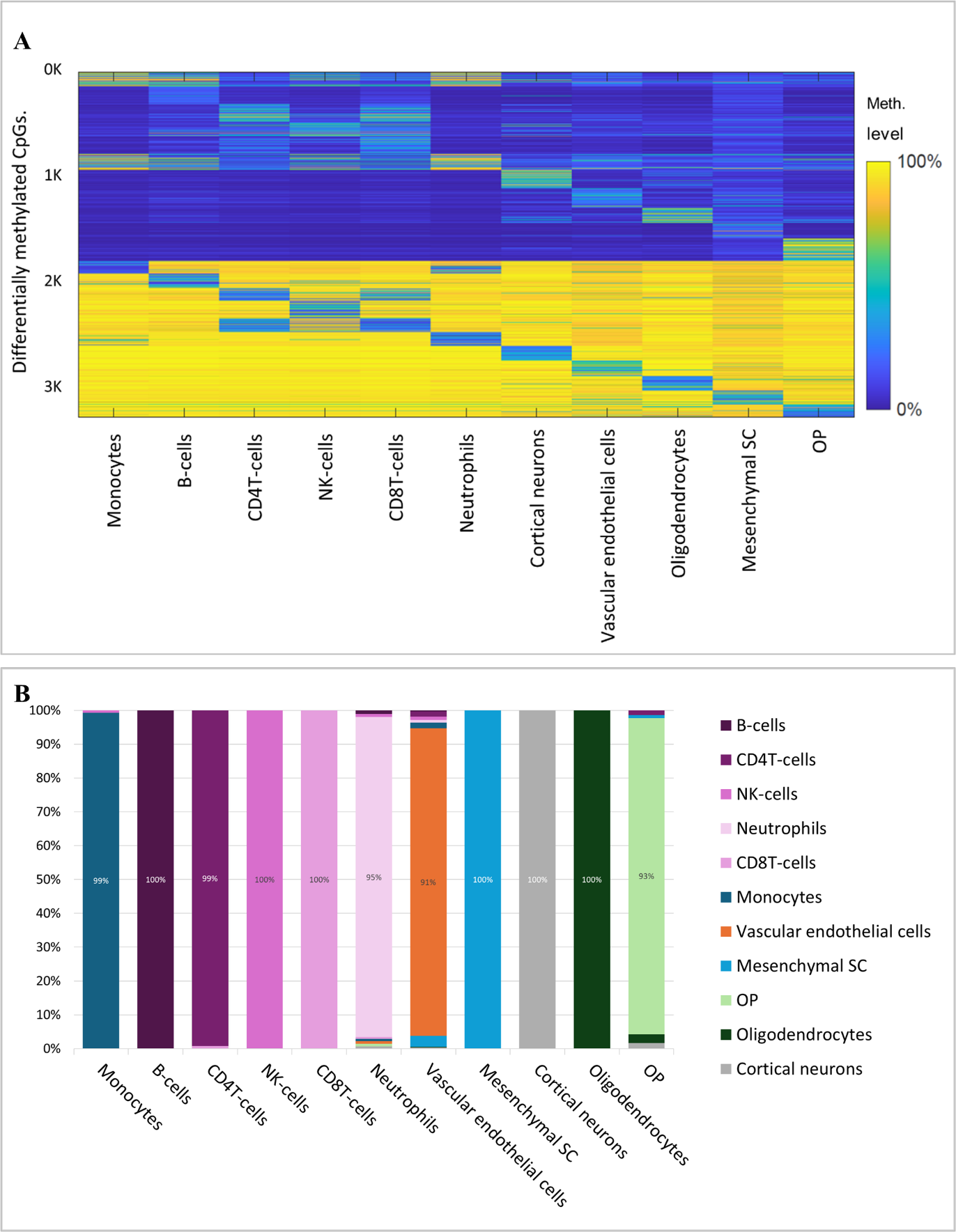

Figure S2: Methylation-based deconvolution of normal cell type samples

(A) The reference atlas used for deconvolution includes 11 tissue-specific cell types (columns) across 3,274 CpG sites (rows). (OP, oligodendrocyte-lineage precursors). (B) Proportions of normal cell types derived from the deconvolution of methylation profiles of various normal cell types (n=1 per sample), using a reference atlas derived from other normal samples.

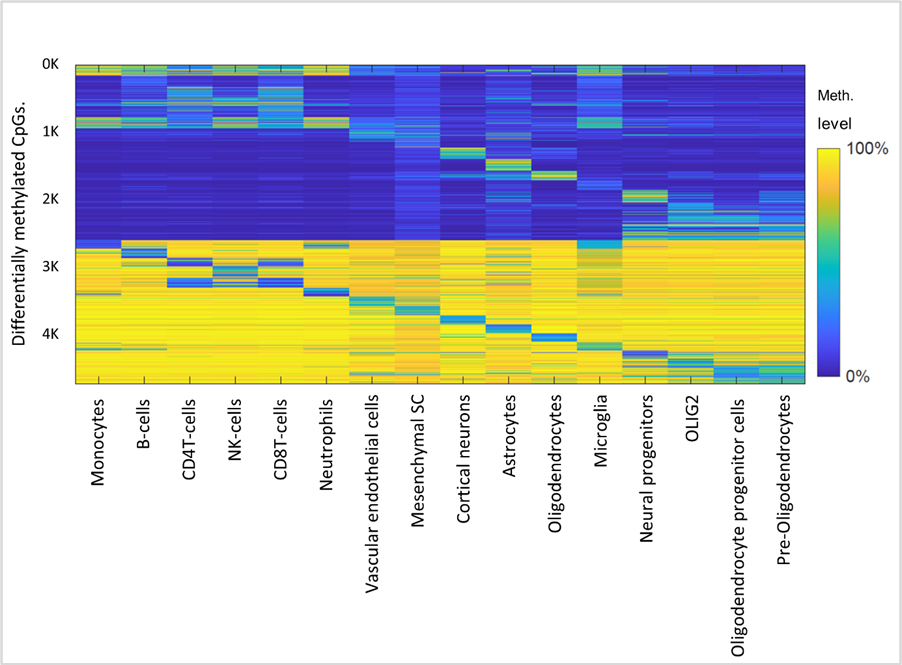

Figure S3. Reference atlas for GBM deconvolution and cell-type identification using methylation signatures.

The methylation atlas comprises 16 tissues/cell types (columns) across 4712 CpGs sites (rows) that are located in 2817 genomic blocks of 500 bp each. Feature selection was carried out as described in a previous study.^20^ Specifically, the top 100 uniquely hypermethylated CpGs and hypomethylated CpGs for each cell type, yielding a total of 3200 tissue-specific CpGs, with neighboring CpGs within 50 bp were also included.

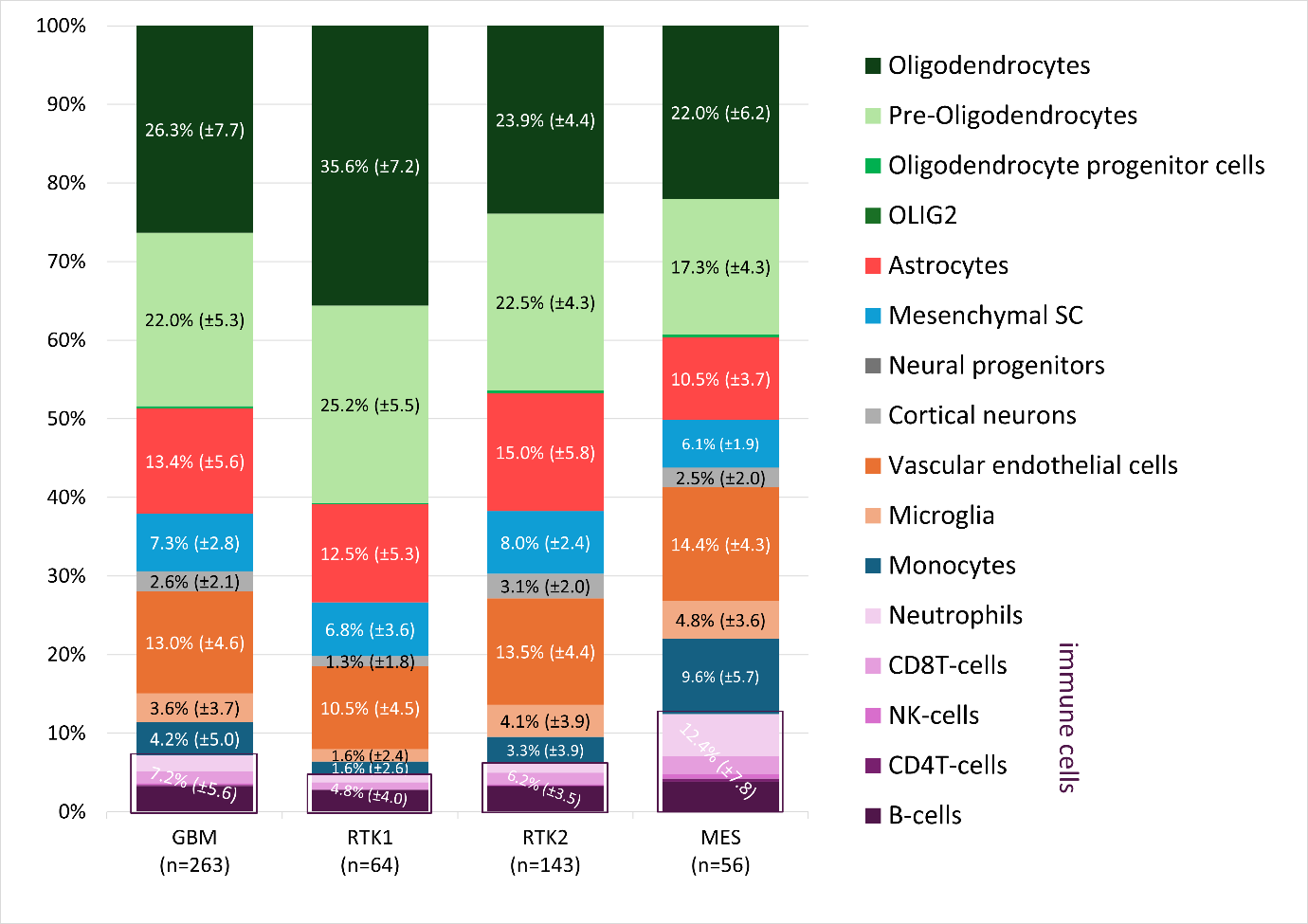

Figure S4. Cell-type proportions derived from deconvolution analysis in GBM and their differences across GBM subtypes, based on atlas of 16 cell types.

Bar plots comparing the average proportions of normal cell types derived from the deconvolution of 263 GBM samples (leftmost bar), subdivided into RTK1 (n=64), RTK2 (n=143), and MES (n=56), based on the atlas of 16 cell types. For each cell type, the bar displays the average percentage along with its standard deviation. The immune cell proportion represents the combined total percentage of all five immune cell components (B cells, CD4 T cells, NK cells, CD8 T cells, and neutrophils), are boxed and shown in purple shading. NP, Neural progenitors; OLIG2, OLIG2-positive progenitor cells; OPC, oligodendrocyte progenitor cells; pre-oligodendrocytes, O4+ pre-oligodendrocytes. The average percentage and standard deviation for each cell type are displayed within its respective bar.

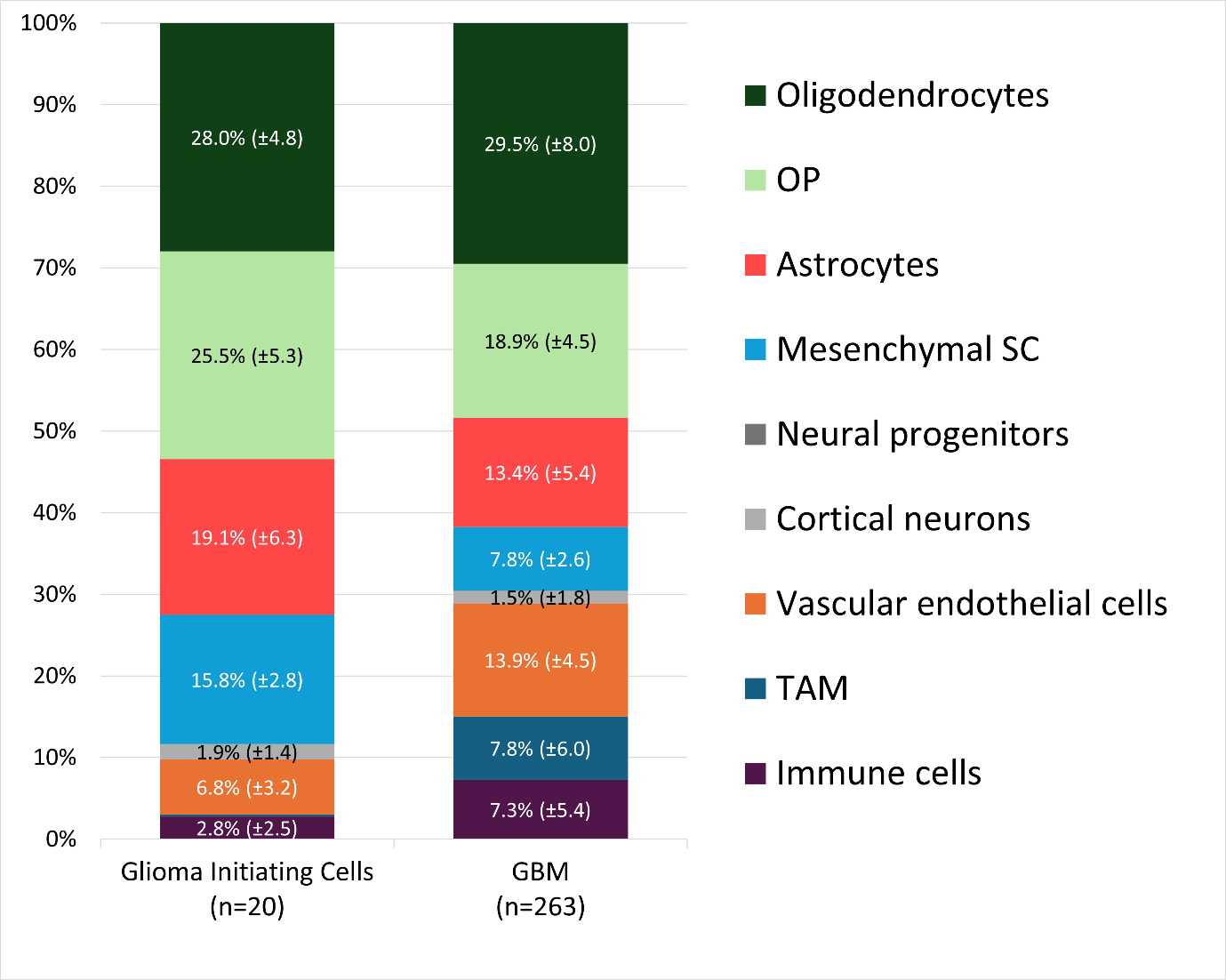
Figure S5: Methylation-based deconvolution of glioma initiating cells (GIC) samples compered to GBM bulk tumors

Bar plots comparing the average proportions of normal cell types derived from the deconvolution of methylation profiles in GBM-derived GIC samples (n=20, left) and bulk of GBM tumors (n=263, right). The average percentage and standard deviation for each cell type are displayed within its respective bar. (OP: oligodendrocyte-lineage precursors).

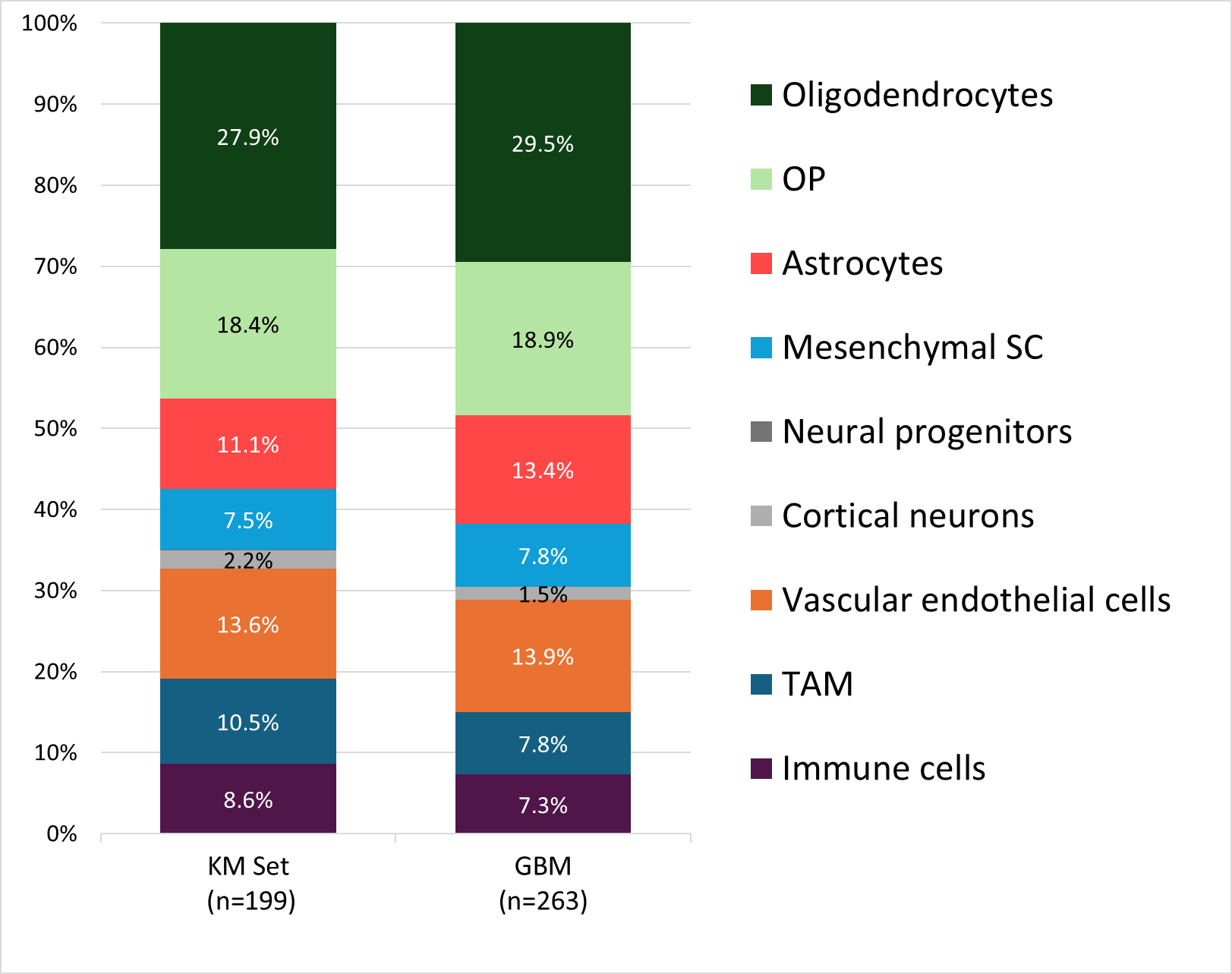
Figure S6: Methylation-based deconvolution of survival analysis set samples compared to GBM bulk tumors.

Bar plots comparing the average proportions of normal cell types derived from the deconvolution of methylation profiles in GBM sample set (n=199, left) collected from TCGA and GEO datasets for Kaplan-Meier (KM Set) survival analysis, and bulk GBM tumors (n=263, right). The comparison of proportion fractions of each cell type showed no significant difference (chi-square goodness-of-fit test, p=0.96). The average percentage is displayed within its respective bar. (OP: oligodendrocyte-lineage precursors).

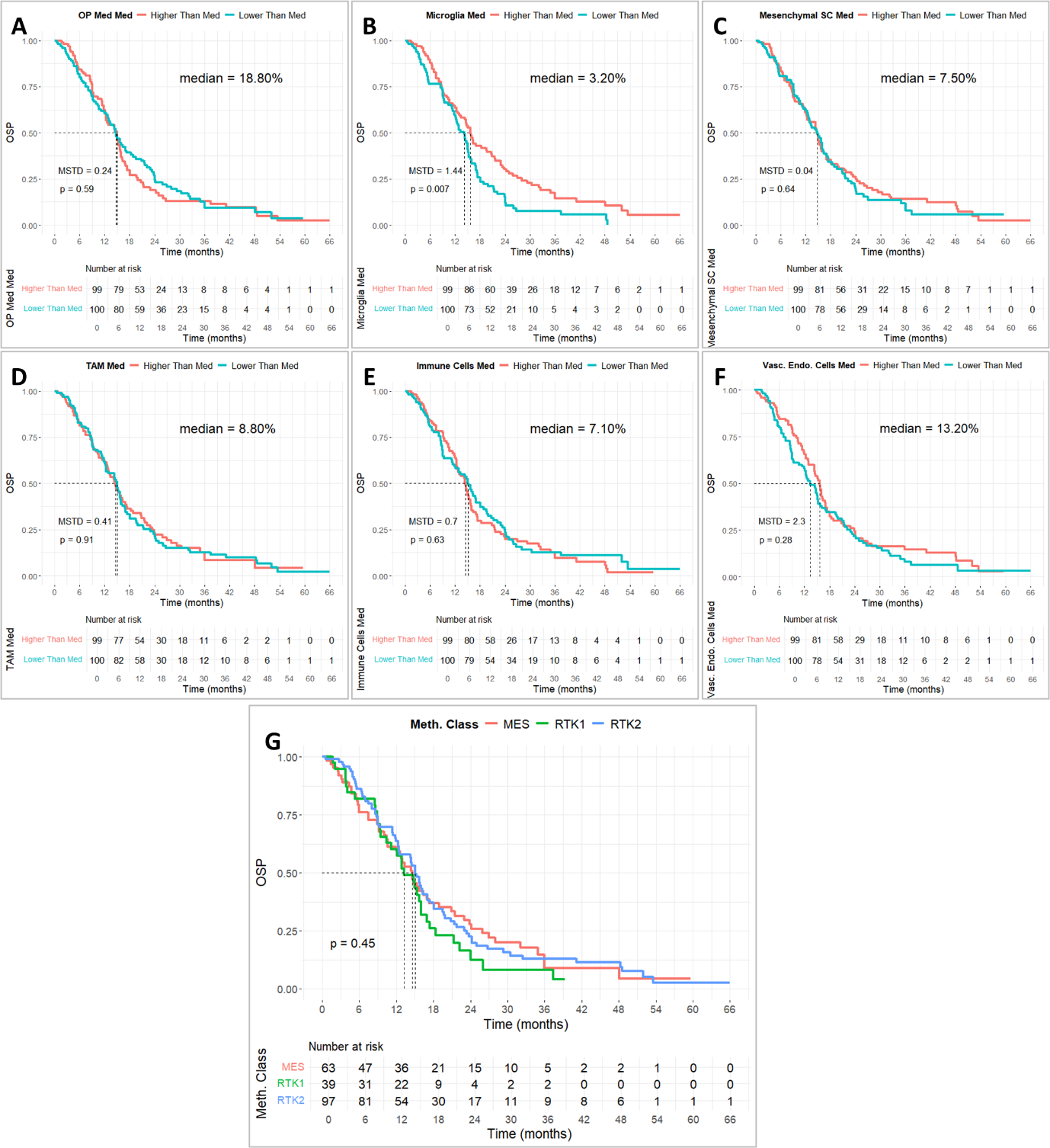

Figure S7: Survival analysis based on cell type proportions and GBM subclasses

(A-F) Kaplan-Meier survival analysis of 199 cases showing differences in overall survival probability based on cell type proportions. The median survival time difference (MSTD) between the groups is indicated in months. The groups are stratified into high and low groups based on median value (High > median, Low < median): (A) oligodendrocyte precursors (OP), (B) microglia, (C) mesenchymal stem cells, (D) TAM (total of monocytes and microglia), (E) immune cells (total of B cells, CD4+ T cells, NK cells, CD8+ T cells, and neutrophils), (F) vascular endothelial cells. (G) Kaplan-Meier survival analysis of 199 GBM samples classified as RTK1 (n=39), RTK2 (n=97), and MES (n=63).

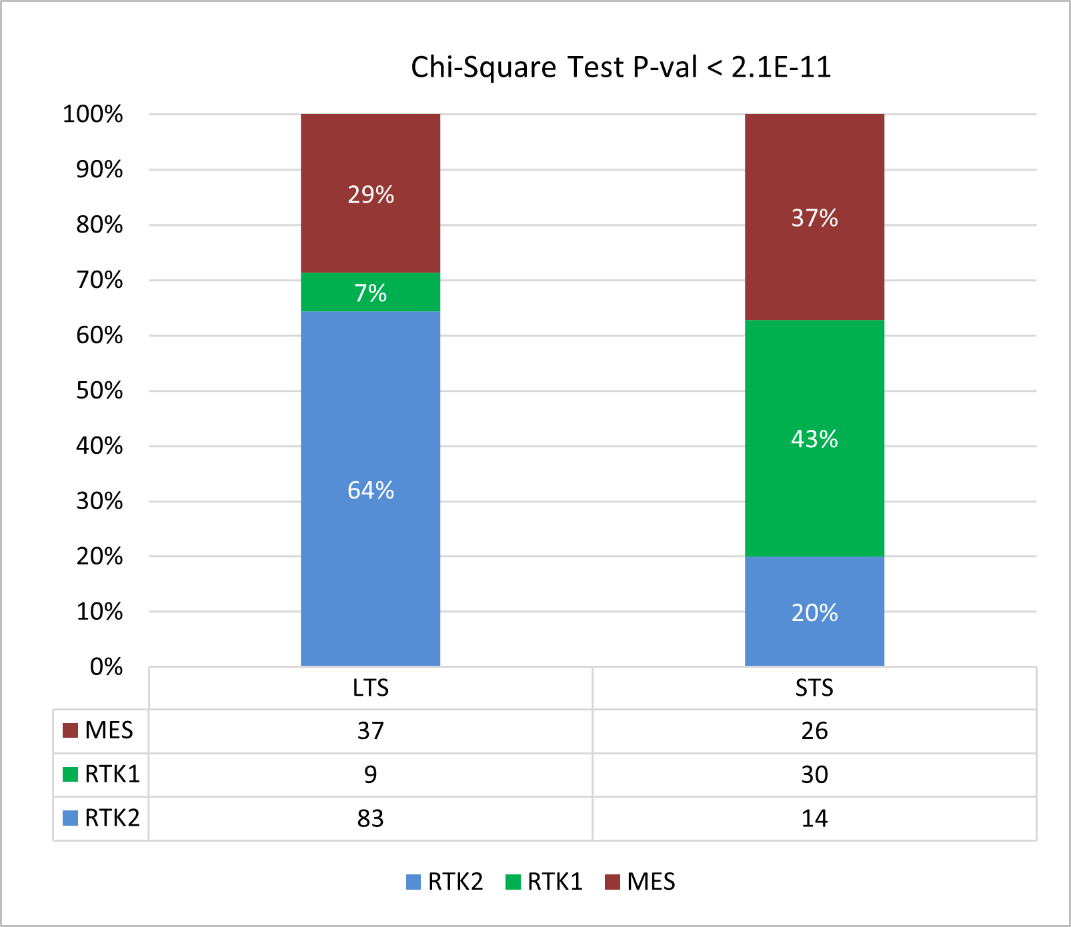

Figure S8: Distribution of GBM subclasses according to the astrocyte-to-oligodendrocyte (Astro/Olig) ratio groups in relation to survival time.

Histogram showing the significant proportional distribution of the three GBM subclasses, RTK1 (n=39), RTK2 (n=97), and MES (n=63), across the two survival groups, long-term survival (LTS) and short-term survival (STS), determined by the astrocyte-to-oligodendrocyte proportion cutoff ratio (chi-square test, p<2.1E-11). The table below provides patient proportions for each group.
